# How well do crop modeling groups predict wheat phenology, given calibration data from the target population?

**DOI:** 10.1101/708578

**Authors:** Daniel Wallach, Taru Palosuo, Peter Thorburn, Emmanuelle Gourdain, Senthold Asseng, Bruno Basso, Samuel Buis, Neil Crout, Camilla Dibari, Benjamin Dumont, Roberto Ferrise, Thomas Gaiser, Cécile Garcia, Sebastian Gayler, Afshin Ghahramani, Zvi Hochman, Steven Hoek, Heidi Horan, Gerrit Hoogenboom, Mingxia Huang, Mohamed Jabloun, Qi Jing, Eric Justes, Kurt Christian Kersebaum, Anne Klosterhalfen, Marie Launay, Qunying Luo, Bernardo Maestrini, Henrike Mielenz, Marco Moriondo, Hasti Nariman Zadeh, Jørgen Eivind Olesen, Arne Poyda, Eckart Priesack, Johannes Wilhelmus Maria Pullens, Budong Qian, Niels Schütze, Vakhtang Shelia, Amir Souissi, Xenia Specka, Amit Kumar Srivastava, Tommaso Stella, Thilo Streck, Giacomo Trombi, Evelyn Wallor, Jing Wang, Tobias K.D. Weber, Lutz Weihermüller, Allard de Wit, Thomas Wöhling, Liujun Xiao, Chuang Zhao, Yan Zhu, Sabine J. Seidel

## Abstract

Predicting phenology is essential for adapting varieties to different environmental conditions and for crop management. Therefore, it is important to evaluate how well different crop modeling groups can predict phenology. Multiple evaluation studies have been previously published, but it is still difficult to generalize the findings from such studies since they often test some specific aspect of extrapolation to new conditions, or do not test on data that is truly independent of the data used for calibration. In this study, we analyzed the prediction of wheat phenology in Northern France under observed weather and current management, which is a problem of practical importance for wheat management. The results of 27 modeling groups are evaluated, where modeling group encompasses model structure, i.e. the model equations, the calibration method and the values of those parameters not affected by calibration. The data for calibration and evaluation are sampled from the same target population, thus extrapolation is limited. The calibration and evaluation data have neither year nor site in common, to guarantee rigorous evaluation of prediction for new weather and sites. The best modeling groups, and also the mean and median of the simulations, have a mean absolute error (MAE) of about 3 days, which is comparable to the measurement error. Almost all models do better than using average number of days or average sum of degree days to predict phenology. On the other hand, there are important differences between modeling groups, due to model structural differences and to differences between groups using the same model structure, which emphasizes that model structure alone does not completely determine prediction accuracy. In addition to providing information for our specific environments and varieties, these results are a useful contribution to a knowledge base of how well modeling groups can predict phenology, when provided with calibration data from the target population.

## Introduction

Crop models are widely used to simulate the effects of weather, soil, and crop management on crop growth and development (Rauff & Bello, 2015; van Ittersum et al., 2003). Here, we focus specifically on the use of crop models to simulate crop phenology, i.e. the cycle of biological events in plants. Matching the phenology of crop varieties to the climate in which they grow is a critical crop production strategy (Hunt et al., 2019; Rezaei et al., 2018, 2015). Accurate simulation of phenology is essential for assessment of the impact of weather conditions on crop growth and development (Ceglar et al., 2019). Process-based models similar to those for crops can be used for natural vegetation, so crop models can serve as examples for studies of phenology in all ecosystem types (Piao et al., 2019).

Given the importance of crop models for predicting phenology, an important question is the accuracy of such predictions. In this study we use prediction in the sense of determining outputs (dates of development stages) from known inputs (weather, soil, management), rather than predicting future events with unknown weather, although, the use of seasonal forecasts for predicting future weather and phenology has also been explored (Canal et al., 2017). There have been multiple studies evaluating how well crop models predict phenology, but results depend on multiple factors, so it is hard to generalize for new situations.

The general practice for evaluation is to use a subset of the available data to calibrate a model, in order to adapt it to a new cultivar or range of environments, and then to evaluate the model using other data not used for calibration. There are, however, important differences between the evaluation studies reported. A major difference between evaluation studies concerns the relation between the calibration and evaluation data. On the one hand, there are studies which intentionally have different ranges of environmental conditions for the calibration and evaluation data. For example, Biernath et al. (2011) calibrated four models using data from well-watered treatments with ambient CO_2_, and subsequently evaluated how well the models simulated results for enhanced CO_2_ treatments with or without water limitations. Hussain et al. (2018) used data from a wheat experiment that included a range of planting dates, and therefore, crop stresses. They used data from the least stressed treatment in the calibration process and evaluated the resultant model on the remaining planting dates at the same location. The evaluation data thus represented a different range of conditions, namely more-stressed conditions, than the calibration data. In a multi-model ensemble study of the effect of high temperatures on wheat growth Asseng et al. (2015) provided modeling groups with detailed crop measurements for one sowing date and the models were evaluated using different sowing dates, some with additional artificial heating. Again, the evaluation data represented a much larger range of temperatures than represented in the calibration data. In such studies one is evaluating, at least partially, aspects of the model for which the calibration data are not pertinent. One could refer to such studies as “extrapolation” studies, since the goal is to evaluate the capability of the model to extrapolate to a range of environments outside those sampled for the calibration data. While the capacity of crop models to extrapolate to conditions quite different from those of the calibration data is obviously of interest, a difficulty with such studies is that the results will depend on how exactly the calibration and evaluation data sets differ.

On the other hand, in the second type of evaluation study, the calibration and evaluation data are drawn from the same underlying “target population” (i.e. range of environments and management for which one wants predictions). One could refer to such studies as “interpolation” studies. (Of course there is sampling variability, so the ranges of conditions in the calibration and evaluation data may not be strictly identical). An example of this type of evaluation is the study by Gouache et al. (2012) who use a modified version of the ARCWHEAT model to predict phenology of the wheat cultivar Soissons in France. In this case, the calibration and evaluation used the same data. Ceglar et al. (2019) used the WOFOST model in a gridded study on wheat throughout Europe and the calibration and evaluation data were sampled from the same population. For crop management, it is often this type of evaluation that is of interest. Based on a sample of data from current conditions and practices, the intention is to evaluate how well the model predicts for other occurrences from the same or a similar range of conditions and practices.

Another difficulty in drawing conclusions from previous evaluation studies relates to the way the evaluation was performed. In conducting an evaluation, the data used for evaluation should be independent of the data used for calibration. If the same data are used for both calibration and evaluation, or more generally if the predictions are correlated with the evaluation data, the estimated error in general underestimates the true error for new environments (Efron, 1986) This is shown explicitly in Ceglar et al. (2019), who found a weighted root mean squared error of about eight days to anthesis for the calibration data, compared to about 11 days for the evaluation data. For days to maturity, the values were about 10 days for the calibration compared to 19 days for the evaluation data. In many evaluation studies, this independence of calibration and evaluation data is not strictly satisfied, for example because the same sites appear in both data sets (Andarzian et al., 2015; Hussain et al., 2018; Yuan et al., 2017).

Overall, it is difficult to generalize from existing evaluation studies, which often involve one or only a few models, or test some specific aspect of extrapolation to new conditions, or do not test on data truly independent of the data used for calibration. Therefore, it would be useful to build up a knowledge base with evaluation studies that do not have those limitations.

In this study, we are specifically interested in wheat phenology in France. For wheat management, predicting growth stages is of major interest as it allows anticipating important management decisions including nitrogen fertilization, irrigation, and crop protection (Bogard et al., 2015). There are multiple decision support tools for wheat farmers in France, including tools commercialized by the French extension institute Arvalis-Institut du vegetal (Le Bris et al., 2015) or the Déciblé tool (Chatelin et al., 2005). In these tools, phenology plays a crucial role. Of particular interest are the date of stem elongation, when the crop nitrogen (N) requirements increase rapidly, and the heading date, which is usually the last date for N fertilization (Bogard et al., 2015).

The purpose of this study was to answer the question how well crop modeling groups can predict wheat phenology in France for current conditions and management, where the calibration data are sampled from the target population. This is thus an example of an interpolation type evaluation. The simulations are evaluated for sites and years not present in the calibration data, meaning that the current study is a rigorous test of how well the simulations can predict for unseen locations and weather conditions. Beyond the specific target population considered here, this study can contribute more generally to a knowledge base of how well crop modeling groups can predict phenology when calibration data are sampled from the target population.

## Materials and Methods

### Experimental data

The crop and climatic data were provided by ARVALIS – Institut du vegetal, a French agricultural technical institute, who run multi-year multi-purpose trials at multiple locations across France, which include variety trials. The data used are from the two winter wheat varieties, namely Apache which is a common variety grown throughout France and Central Europe and Bermude, mainly grown in Northern and Central France. The trials have three repetitions and follow standard agricultural practices, with N fertilization sufficient to avoid N stress. Weather data are from weather stations near each field. The target population from which both the calibration and evaluation data are samples is sites in Northern France where winter wheat is grown (Figure 1), under current conditions and subject to standard management.

**Figure 1.**
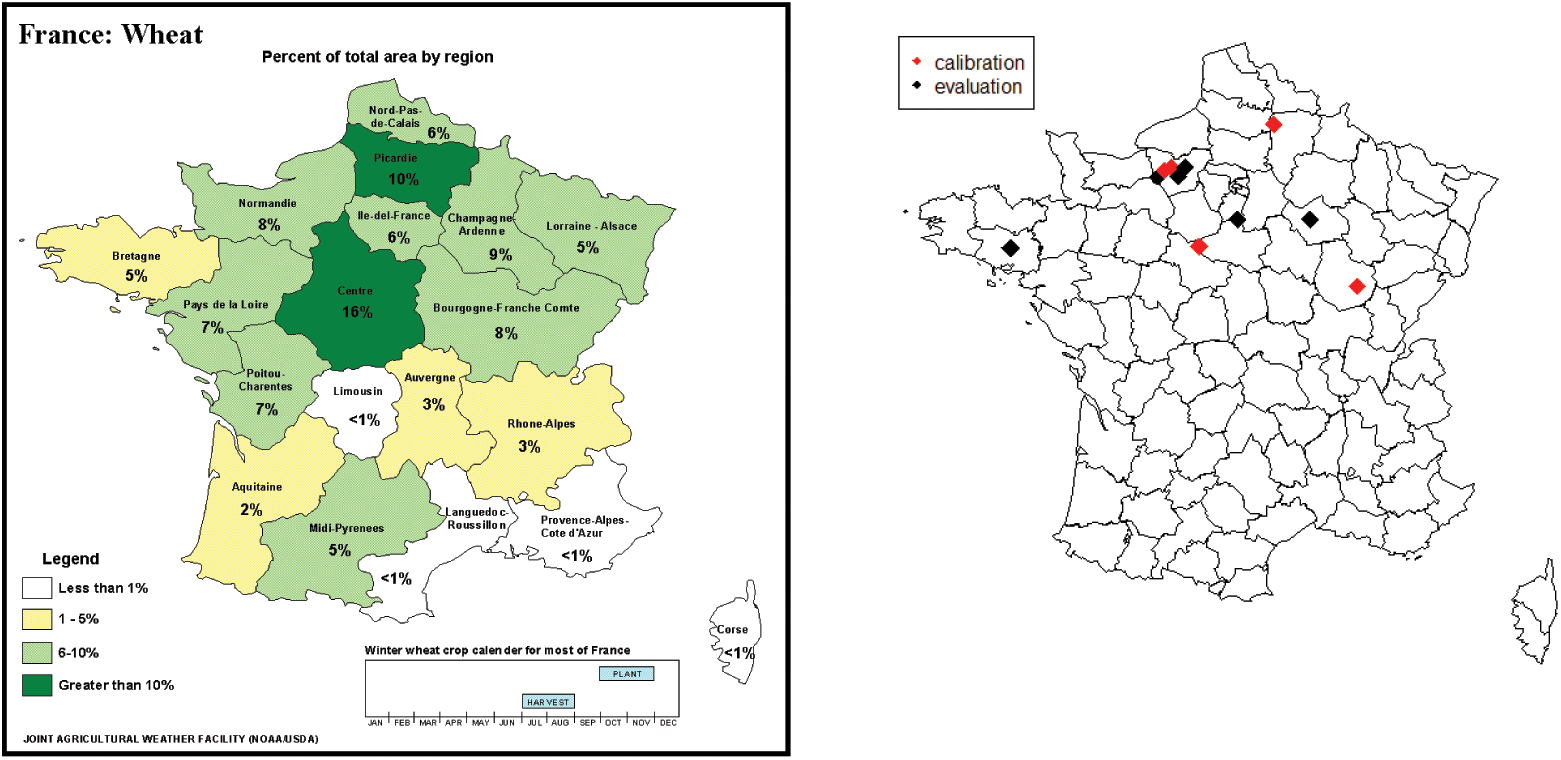
Major wheat growing regions in France (left) and locations of sites that provided data (right).

The observed data used for model calibration and evaluation are the dates of two development stages, namely beginning of stem elongation (growth stage 30 on the BBCH and Zadoks scales) (Lancashire et al., 1991; Zadoks, Chzang, & Konzak, 1974)) and middle of heading (growth stage 55 on the BBCH and Zadoks scales). These two stages are of practical importance because they can easily be determined visually and are closely related to the recommended dates for the second and third N fertilizer applications. For BBCH30, observations were made on multiple days around stage BBCH30, and then a simple correction model was used to determine the exact date when the stem was 1 cm long. It is estimated that the uncertainty is 3-4 days for determination of this stage. The middle of heading is easier to observe. The estimated uncertainty of the observed value for this stage is about one day.

To characterize the environments (Figure 2a and Supplementary Figure S1, we calculated average temperature from sowing to BBCH30 and to BBCH55, average photoperiod from sowing to BBCH55 calculated using the daylength function in the R package insol (Corripio, 2019.; R Core Team, 2017) and days to full vernalization, calculated using the model in van Bussel, Stehfest, Siebert, Müller, & Ewert (2015) with the value *V*_*sat*_ = 50 days, estimated from the figure in their paper.

**Figure 2.**
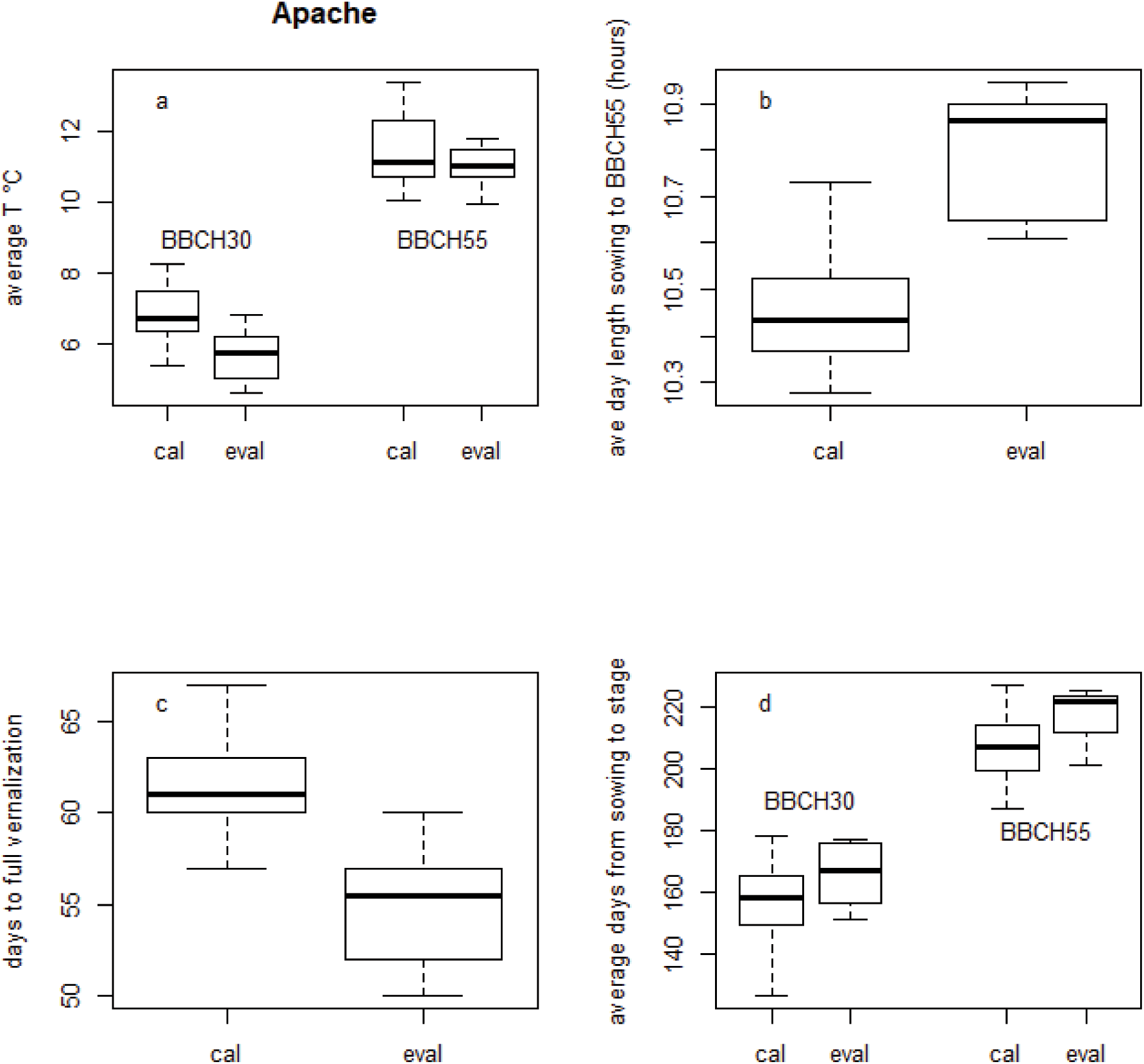
Characteristics of calibration (“cal”) and evaluation (“eval”) data sets. Average daily temperature from sowing to BBCH30 and from BBCH30 to BBCH55 (a). Average day length from sowing to BBCH30 (b). Days to vernalization calculated as explained in text (c). Average days from sowing to observed BBCH30 and to observed BBCH55 (d).

The data were divided into two parts (Table 1). For one set of environments, the calibration environments (six sites, five years for a total of 14 environments i.e. site-year combinations), the observed days of BBCH30 and BBCH55 were provided to participating modeling groups for calibration. For the second set of environments, the evaluation environments (five sites, two years for a total of eight environments), weather and management were provided but not observed days of BBCH30 and BBC55. The division of the data was such that the calibration and evaluation data had no sites or years in common.

**Table 1.** Environments (site-year combinations) that provided the data. C= calibration environments. E = evaluation environments. Dates shown are sowing dates (in year before harvest year), which are the same for both varieties.

| Site<br>(longitude,latitude) | Harvest year |  |  |  |  |  |  |
| --- | --- | --- | --- | --- | --- | --- | --- |
|  | 2010 | 2011 | 2012 | 2013 | 2014 | 2015 | 2016 |
| FORESTE<br>(3.20,49.82) |  |  | E<br>Oct 11 | E<br>Oct 23 |  |  |  |
| MERY<br>4.02,48.33) | C<br>Oct 8 | C<br>Oct 7 |  |  | C<br>Oct 7 | C<br>Oct 15 |  |
| ROUVRES<br>5.09,47.28) |  |  | E<br>Oct 11 | E<br>Oct 18 |  |  |  |
| CESSEVILLE<sup>1</sup><br>(0.90,49.15) |  | C<br>Oct 13 |  |  |  |  |  |
| IVILLE<sup>1</sup><br>(0.90,49.15) |  |  | E<br>Oct 12 |  |  |  |  |
| VILLETES<sup>1</sup><br>(0.90,49.15) |  |  |  | E<br>Oct 24 |  |  |  |
| EPREVILLE<sup>1</sup><br>(0.90,49.15) |  |  |  |  | C<br>Oct 15 |  |  |
| CRESTOT<sup>1</sup><br>(0.90,49.15) |  |  |  |  |  | C<br>Oct 14 |  |
| OUZOUER<br>(1.52,47.90) |  |  | E<br>Oct 21 | E<br>Oct 30 |  |  |  |
| BIGNAN<br>(-2.73,47.88) | C<br>Oct 30 | C<br>Oct 28 |  |  | C<br>Oct 21 | C<br>Oct 27 |  |
| BOIGNEVILLE<br>(2.38,48.33) |  |  |  |  | C<br>Oct 21 | C<br>Oct 23 | C<br>Oct 21 |

<sup>1</sup> These are separate sites that are geographically close to one another and share a single weather station.

The background and input information provided to the participating modeling groups for all environments included information about the sites (location, soil texture, field capacity, wilting point), management (sowing dates, sowing density, irrigation and fertilization dates and amounts), and daily weather data (precipitation, minimum and maximum air temperature, global radiation, and potential evapotranspiration). Initial soil water and N content were not measured in these experiments, instead best estimates were provided by the experimental scientists. If any models required other input data, modeling groups were asked to derive those values themselves in a way that seemed appropriate.

### Simulated values

Twenty-seven modeling groups participated in this study, noted M1-M27. We distinguish here between model structure, i.e. the mathematical form of the model equations, which is associated with a model name, and modeling group. The modeling group chooses not only a specific model structure, but also the calibration approach and the values of the fixed parameters, i.e. those not changed by calibration. References for the model structures are given in Supplementary table S1. The four groups M2, M3, M4, and M5 all used the same model structure (i.e. models with the same name), noted model structure S1. The four models M7, M12, M13, and M25 also shared a common model structure, noted S2, and groups M23 and M24 both used model structure noted S3. In the presentation of the results the modeling groups are anonymized and are identified simply as M1 to M27. It would be misleading to use the names of the model structures, since different groups using the same model structure obtained different results.

Two of the modeling groups (M9, M18) only simulated days to development stage BBCH55 and not to stage BBCH30. Results for these two models are systematically included with the results for the other models, but averages over development stages for these two models only refer to BBCH55. This is not repeated explicitly every time an average over development stages is discussed.

In addition to the results of the individual modeling groups, we define two ensemble predictors. The model that predicts phenology using the mean of the simulated values is termed “e-mean” and the model that predicts using the median of the simulated values is termed “e-median”.

We also define two very simple predictors, as a basis for skill measures. The first, denoted the “naive” model, predicts phenology using the average of the observed values in the calibration data. The naive model predictions for days from sowing to BBCH30 and BBCH55 are respectively 155.9 and 206.9 days for variety Apache, and 156.1 and 213.1 days for variety Bermude. The second simple predictor, the “onlyT” model, assumes that the sum of degree days (DD) to BBCH30 or BBCH55 will always be the same as in the calibration data. A base temperature of 0°C, is assumed and so DD on any day is equal to the mean temperature that day. The onlyT model predicts that BBCH30 and BBCH55 will occur 1036 and 1620 DD after sowing respectively for variety Apache, and 1039 and 1701 DD after sowing for variety Bermude. The use of 0°C as base temperature is simple, but the actual base temperature for development may differ between varieties and be in general somewhat higher (Salazar-Gutierrez et al., 2013).

### Calibration and simulation experiment

The participants were provided with observed phenology data (dates of BBCH30 and BBCH55) only for the calibration environments. The participants were asked to calibrate their model using those data, and then to use the calibrated model to simulate phenology for both the calibration and evaluation environments. No guidelines for calibration were provided. Participants were instructed to calibrate their model in their “usual way”. At no time did the participants see the evaluation data.

### Evaluation

Our basic measure of error is the mean absolute error (MAE). This metric is not as widely used as mean squared error (MSE) or root mean squared error (RMSE) but is preferred here as a more direct measure of error (Willmott and Matsuura, 2005). MAE for modeling group *m*, for predicting days from sowing to either stage, is:

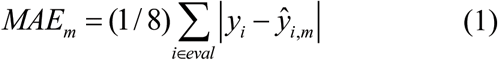

where the sum is either over the eight environments used for evaluation or over the 14 environments of the calibration data, and *y*_*i*_ and *ŷ*_*i,m*_ are respectively the observed and simulated values for one of the two varieties. The corresponding expressions for MSE and RMSE are

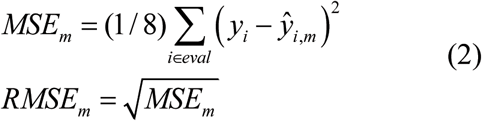

We also calculate two skill scores, based on the models naive and onlyT. The skill score EF compares MSE for a modeling group to MSE for the naive model:

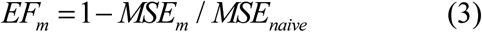

where an EF = 1 indicates a modeling group providing perfect predictions and an EF <= 0 indicates a modeling group that does no better than the naive model. It is important to emphasize that the naive model here is based only on the calibration data, so it is a predictor that could be used in practice. The second skill score, skillT, has the same form as EF, but compares MSE of a modeling group to MSE for the onlyT model rather than to MSE for the naive model:

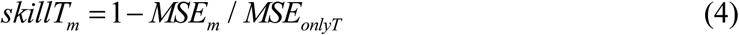

### Within- and between-structure variability

Since three model structures are represented by multiple modeling groups, it is possible to estimate separately within-structure and between-structure variability. We use the law of total variance (Casella and Berger, 1990):

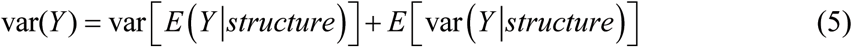

where *Y* is either days from sowing to BBCH30 or to BBCH5. The first term is the variance between structures, where each structure is represented by the expectation over groups using that structure. This is the between-structure contribution to total variance, and is estimated by first calculating the mean simulated value for each model structure, and then the variance between structures. For structures represented by a single modeling group, the mean is simply the simulated value for that group. The second term is the within-structure contribution. It is estimated based on those structures represented by more than one group. The variance of the simulated values for each of those structures is calculated, and then those variances are averaged.

## Results

### Characteristics of calibration and evaluation data

Both the calibration and the evaluation data are sampled from fields and years in the major wheat growing region of France with standard management and current climate. Nevertheless, sampling variability results in ranges of characteristics that are somewhat different in the two data sets (Figure 2, Supplemenary Figure S1). Differences in median values between the calibration and evaluation data sets are very similar for Apache and Bermude. Differences are approximately 1°C for average temperature to BBCH30 and approximately 0.3°C for average temperature from BBCH30 to BBCH55, 0.4 hours in average day length, and 5.5 days in days from sowing to full vernalization. Despite these differences, the range of days to each stage is similar for the calibration and evaluation data sets.

### Prediction error

Boxplots of MAE for the evaluation data are shown in Figure 3. Median MAE values (and ranges) are 5 (3-18), 6 (3-13), 6 (4-20), and 6 (2-16) days for Apache stage BBCH30, Apache stage BBCH55, Bermude stage BBCH30 and Bermude stage BBCH55, respectively. When errors are averaged over the four predictions (two varieties, two stages), overall MAE is on average seven days, with a range between modeling groups from three to 13 days (Supplementary Table S2). The corresponding RMSE values are a bit larger, with an average of eight days and a range from four to 15 days.

**Figure 3.**
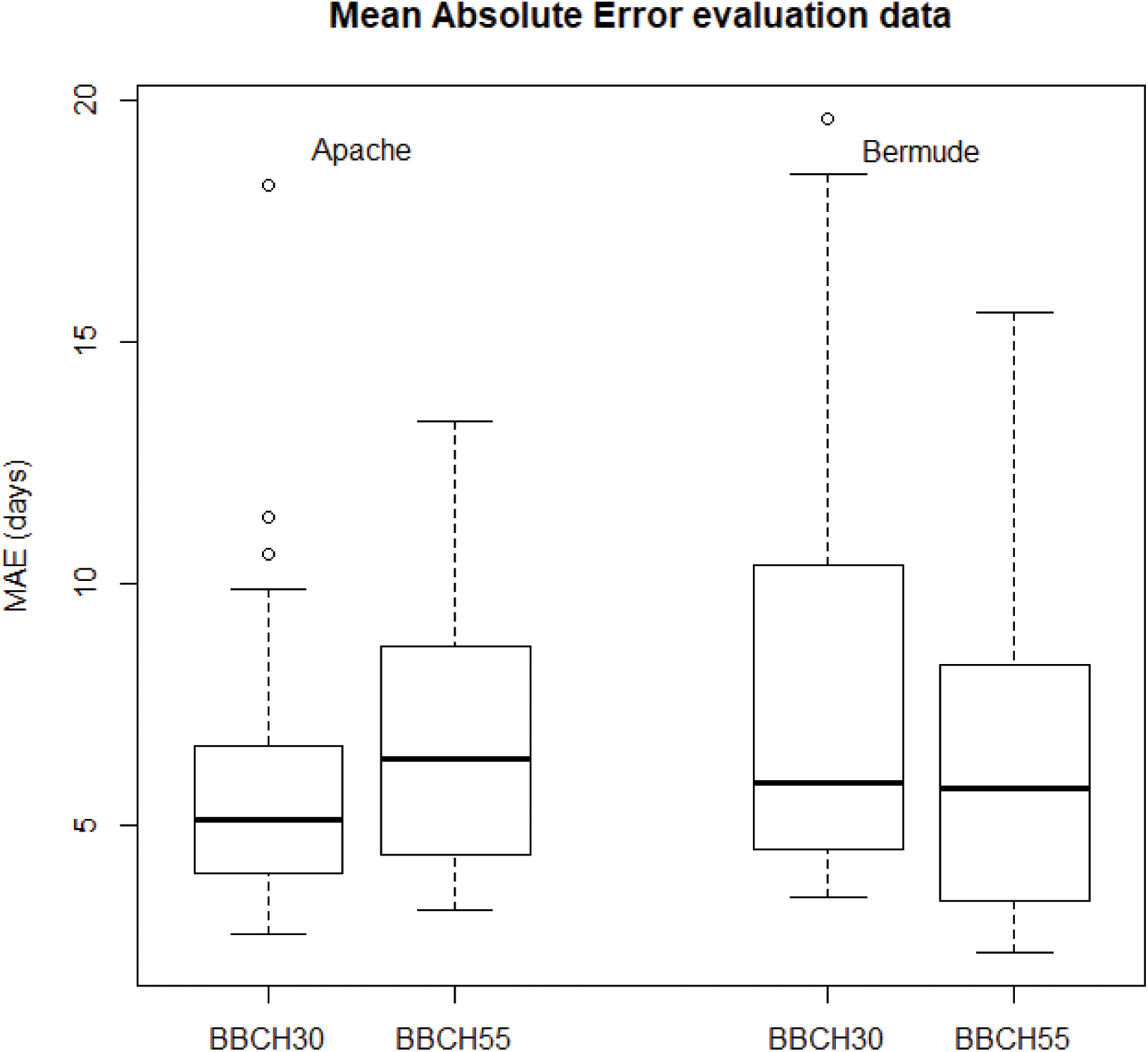
Boxplots of mean absolute error (MAE) for evaluation data, for both varieties and both phenology stages BBCH30 and BBCH50.

The ensemble predictors e-mean and e-median are nearly as good as the best predictor, with e-median slightly better than e-mean. MAE values for e-median are 2, 4, 5, and 3 days for Apache stage BBCH30, Apache stage BBCH55, Bermude stage BBCH30, and Bermude stage BBCH55, respectively.

The naive model has MAE values for the evaluation data of 11 days for BBCH30 for both varieties, and 10 days (Apache) and12 days (Bermude) for BBCH55. The onlyT values are slightly better than those for the naive model for Apache (MAE=10 days for both stages) and slightly worse for Bermude (MAE=13 for days to BBCH30 and MAE=11 for days to BBCH55). Boxplots of the skill measures are shown in Figure 4. Over the 104 predictions by all the modeling groups for the evaluation data, 91 (88%) are better than the naive model (i.e. have EF>0) and 94 (90%) are better than the onlyT model (i.e. skillT>0).

**Figure 4.**
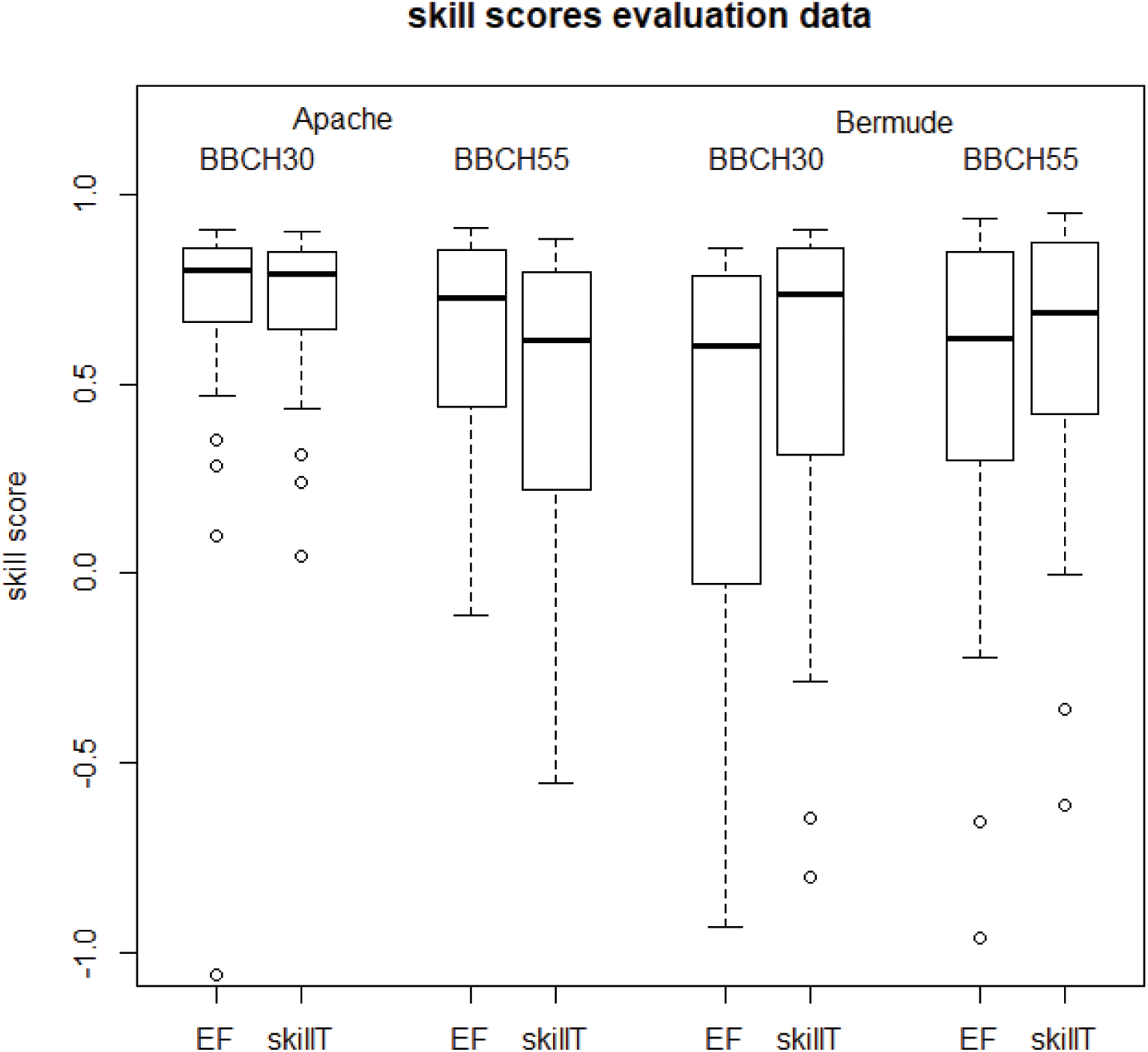
Boxplots of skill measures for the evaluation data.

### Relation of calibration to evaluation errors

MAE values for the calibration data are plotted against the evaluation data in Figure5. The regression is highly significant (*p*<0.001). The means of calibration and evaluation MAE are 5.2 and 6.1 days respectively.

### Differences among modeling groups using the same model structure

Figure 6 shows predictions for each of the modeling groups and for each of the evaluation environments. The simulated values from groups using the same model structure are identified. Averaged over the four predictions, the within- and between-structure standard deviations are respectively 5.6 and 8.0 days, giving a ratio within/between=0.70.

**Figure 5.**
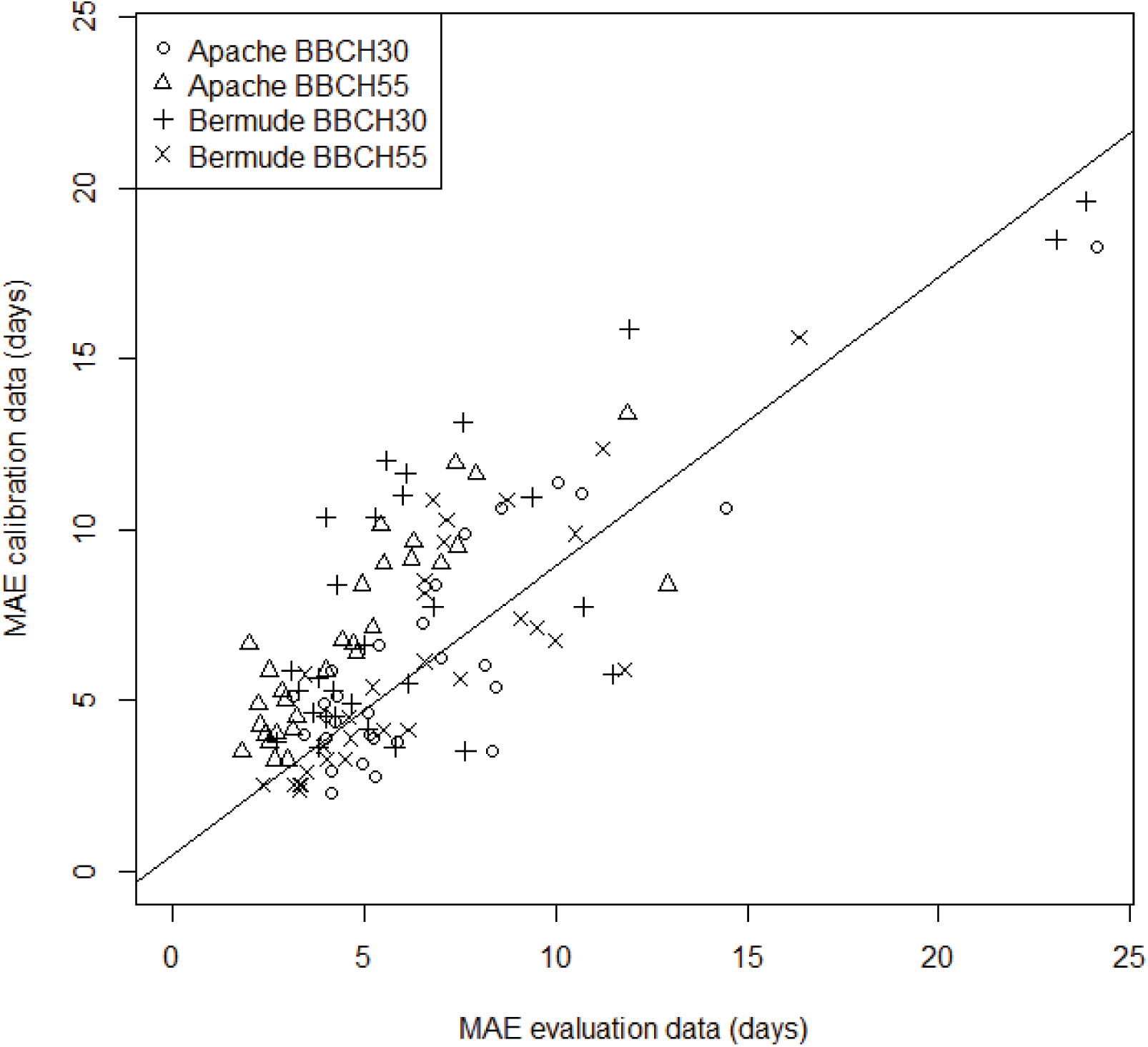
MAE for calibration data versus MAE for the evaluation data. The linear regression line is shown: MAE calibration = 2.42+0.71*MAE evaluation

**Figure 6.**
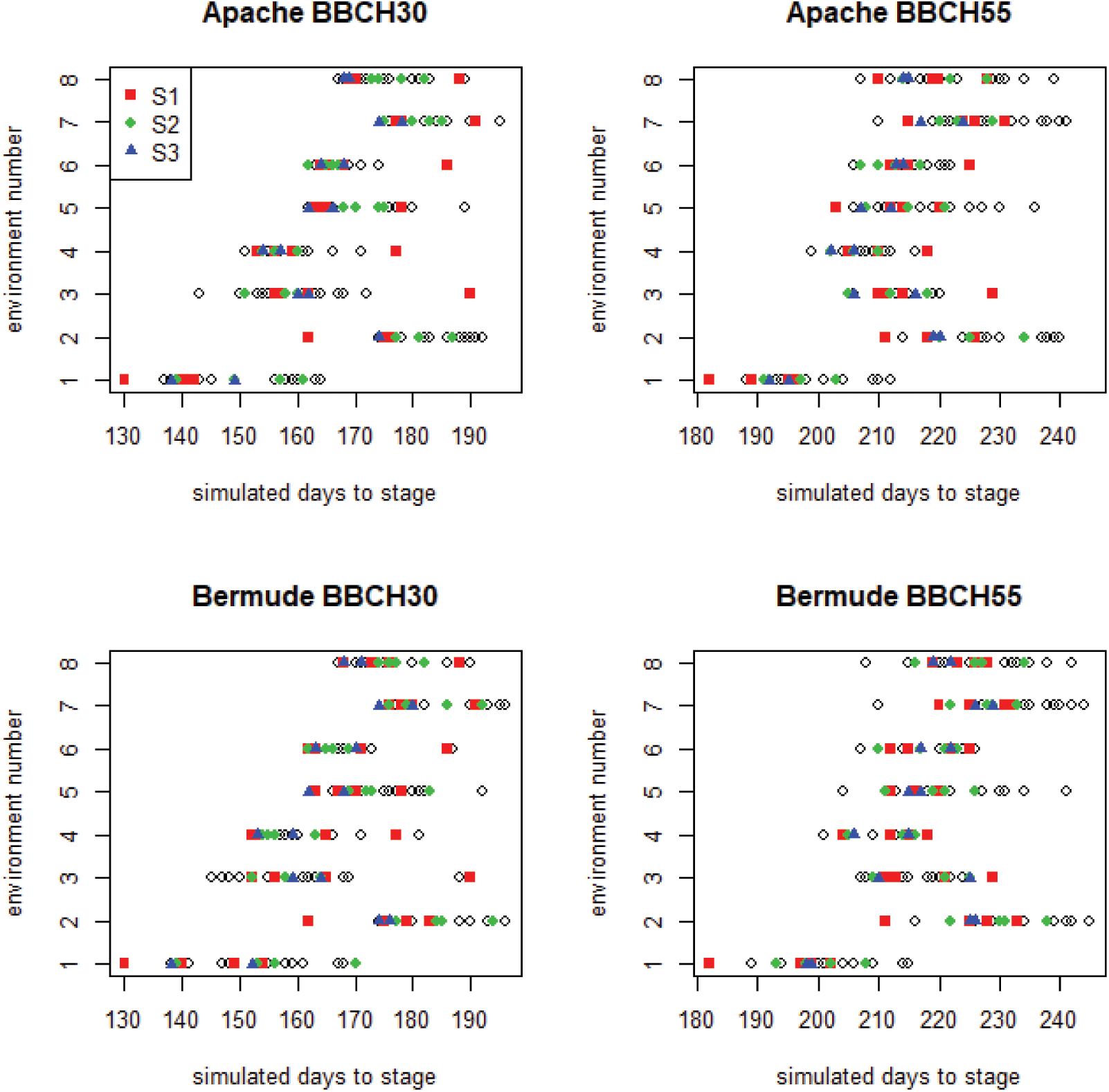
Predictions by each of the original models for each of the evaluation environments. Simulated values from the four modeling groups using model structure S1 are in blue, the four groups using model structure S2 are in red and the two groups using model structure S3 are in green. Other modeling groups are represented by open diamonds.

## Discussion

### Characterization of environments

Even though the calibration and evaluation data are sampled from the same target population, sampling variability leads to some differences between the two sets of environments. This no doubt contributes to prediction error. However, the differences are fairly small. In any case, it is important to characterize environments in order to document the differences between the calibration and evaluation environments and for comparison with other studies.

### Prediction error

The challenge in this study was to predict the time from sowing to beginning of stem elongation and to heading in winter wheat field trials performed across France. This is a problem of practical importance, since these two development stages are important for wheat management, including fertilization and pest control, and the prediction of phenology is also central to several decision support tools (Chatelin et al., 2005; Le Bris et al., 2015).

We consider the specific situation where one has calibration data that is a sample from the target population. This is quite different than tests of how well crop models extrapolate to new conditions that are different than those of the calibration data. Our situation is representative of the practical problem of using a network of trails for calibration, with a view to predicting for similar conditions but at other sites and with other weather. We carry out a rigorous evaluation of how well modeling groups can simulate phenology for sites and years not used for calibration.

Since the differences between the two varieties and between the two stages of interest are relatively slight, we focus on MAE averaged over all four predictions. The best modeling group has overall MAE of 3 days, essentially the same as estimated measurement error for BBCH30, though a bit larger than estimated measurement error of one day for BBCH55. The average over models of overall MAE is seven days, and the average of overall RMSE is eight days.

The prediction errors here are consistent with values reported for individual modeling groups for other European environments. RMSE values of 9 or 8 days were found for two versions of the CERES-wheat model evaluated using a large data set from Germany where calibration and evaluation data were sampled from the same target population (Johnen et al., 2012). Ceglar et al. (2019) used the WOFOST model in a gridded study on wheat throughout Europe. The calibration and evaluation data were sampled from the same population. For Northern France, they found a weighted root mean squared error of about 11 days for days from sowing to anthesis and about 19 days for days to maturity. However, they did not always have true sowing dates, which presumably increased prediction error.

Several studies have evaluated goodness-of-fit (i.e., used the same data for calibration and evaluation) for predicting wheat phenology in Europe. Palosuo et al. (2011) found root mean squared error (RMSE) values of 5 to 20 days for prediction of anthesis date for wheat in Europe by eight different models. Liu et al. (2018) found RMSE values of about 3 days for time to flowering of wheat using the WHEATGROW model. Gouache et al. (2012), using a modified version of the ARCWHEAT model to predict phenology of wheat cultivar Soissons in France, found an RMSE of 4.6 days for stage BBCH30 and 2.5 days for stage BBCH55. These values probably underestimate errors for predicting for new environments, since it is known that fit to calibration data in general is better than prediction for new environments (Efron, 1986). However, the results here suggest that the underestimation may not be too large; the average difference here between MAE values for the calibration and evaluation data is only about one day.

Overall, the range of MAE values found here, from three to 13 days depending on the modeling group (and of RMSE values from four to 15 days), seems to reasonably represent the range of crop phenology prediction error for many European environments. Of particular interest in the results here is the minimum value of MAE of three days, which indicates that phenology can be very well predicted with current models if there is substantial calibration data that is sampled from the target population. Also of particular interest is the average MAE value of seven days, which indicates how well phenology is predicted on the average over a large number of currently active modeling groups. Put another way, one would on average expect MAE of seven days for a modeling group chosen at random, that is without any specific information about prediction error.

Skill measures give a different perspective on prediction error. They indicate whether a model can predict better than some simple alternative. The skill measure EF in this study compares prediction error to that of the naive model, which predicts that days to any stage will be the same as the average days to that stage for the calibration data. Other studies often calculate modeling efficiency by comparing a crop model with average results based on all the data. That average is not a feasible predictor, since it requires knowledge of the results to be predicted. The naive model here, on the other hand, is a feasible predictor since it only requires knowledge of the calibration data. The large majority of modeling groups do better than this simplistic model. Our other simple model, onlyT, based just on degree days, does not do better than the naive model. It seems that vernalization and photoperiod effects are sufficiently important and that temperature by itself is not a good predictor for the two BBCH stages analyzed. Models are even more skillful compared to onlyT than they are compared to naive.

### Ensemble predictors

The above considerations concern the use of the results of a single modeling group. Many other studies have found that the ensemble predictors e-mean and e-median are good predictors (Bassu et al., 2014; Wallach et al., 2018). We similarly find that those ensemble predictors are very nearly as good as the best modeling group.

E-mean and e-median are based on giving equal weight to every simulated value. In other fields there have been suggestions that the results of different modeling groups should be weighted differently, for example, based on how well they fit the calibration data (Haughton et al., 2015). It has also been suggested that differential weighting might be of interest for crop models (Wallach et al., 2016), but this does not yet seem to have been tested. The results here show that errors for the calibration and the evaluation data are very significantly correlated, which supports the idea that it might be worthwhile to predict by giving extra weight to modeling groups that have low error for the calibration data. This merits further investigation. This might be specific to cases where calibration data are drawn from the target population. If the crop model is used for extrapolation, one would expect that the fit to the calibration data is less indicative of the fit to the evaluation data.

### Differences among modeling groups using the same model structure

In this study, three of the model structures were used by more than a single modeling group. This allowed us to estimate separately within-structure and between-structure variability. The within-structure variability, as measured by standard deviation, is about 0.7 times as large as the between-structure standard deviation. The within-structure variability is presumably due to differences in parameter values, which includes differences in values of those parameters not changed by calibration (the fixed parameter values) and the values of the parameters estimated by calibration. Since all modeling groups have the same data for calibration, the differences in calibrated parameters are due to differences in calibration approach and also to differences in the fixed parameters. A study by Confalonieri et al. (2016) found that the effect of the “user” (who is responsible for model set-up and calibration) was larger than the effect of structure, but in that study the users were students with probably less expertise than the modeling groups that participated here.

The emphasis in model evaluation studies is often on the role of model structure, i.e. model equations (Maiorano et al., 2017; Svystun, Bhalerao, & Jönsson, 2019; Wang et al., 2017). Clearly however, as illustrated here, the simulated values depend not only on model structure but also on the parameter values. An important corollary is that model improvement could also be achieved not only by changing equations, but also by improving parameter values, and in particular calibration approach.

## Conclusions

We have provided a rigorous evaluation of how well crop modeling groups can predict phenology of two wheat varieties in Northern France, in the situation where the calibration data are sampled from the target population. This is a situation of practical interest, that arises for instance when one has access to data from a network of variety trials and wants to use the model for management decisions. It is found that the average MAE over all modeling groups is seven days, that the best models and also the mean and median of simulated values, have errors that are comparable to measurement error, but that there are large differences between modeling groups. This suggests that crop models can potentially be useful tools for management decisions for winter wheat in France. The results for two skill scores show that almost all models are better than simplistic alternatives such as using as predictor the average days or average temperature sum to each stage in the calibration data.

It was found that the errors in fitting the calibration data were significantly correlated with errors in prediction. This is not surprising since the calibration data are drawn from the target population. The correlation between goodness-of-fit to the calibration data and prediction accuracy for the evaluation data imply that it is reasonable to choose models, or to choose how to weight different models, based on the calibration results. However, this may not be the case when models are used to predict far outside the range of the calibration data. This emphasizes that it is important to distinguish two types of model evaluation, which can be called “interpolation evaluation” and “extrapolation evaluation”. Here we in an interpolation context.

We found important differences in prediction by different groups using the same model structure. Thus, model inter-comparison studies should be termed “modeling group” inter-comparison studies, since differences depend not only on model structure but also on the values of fixed parameters and the calibration approach chosen by the group. An important implication is that model improvement could result not only from better model structure (i.e. model equations), but also from better parameter values and in particular from better calibration procedures.

## Supporting information

Wallach_et_al_2020_SUPPLEMENTARY_MATERIAL

## Acknowledgements

This work was in part supported by the Collaborative Research Center 1253 CAMPOS (Project 7: Stochastic Modelling Framework), funded by the German Research Foundation (DFG, Grant Agreement SFB 1253/1 2017), the Academy of Finland through projects AI-CropPro (316172) and DivCSA (316215) and Natural Resources Institute Finland (Luke) through a strategic project BoostIA, by the German Federal Ministry of Education and Research (BMBF) in the framework of the funding measure ‘Soil as a Sustainable Resource for the Bioeconomy - BonaRes’, project BonaRes (Module A): BonaRes Center for Soil Research, subproject ‘Sustainable Subsoil Management – Soil^3^’ (grant 031B0151A), the Deutsche Forschungsgemeinschaft (DFG, German Research Foundation) under Germany’s Excellence Strategy - EXC 2070 – 390732324 (PhenoRob), the project BiomassWeb of the GlobeE programme (Grant number: FKZ031A258B) funded by the Federal Ministry of Education and Research (BMBF, Germany), the INRA ACCAF meta-programme, the German Federal Ministry of Education and Research (BMBF) in the framework of the funding measure “Soil as a Sustainable Resource for the Bioeconomy – BonaRes”, project “BonaRes (Module B): BonaRes Centre for Soil Research, subproject B” (grant 031B0511B), the National Key Research and Development Program of China (2017YFD0300205), the National Science Foundation for Distinguished Young Scholars (31725020), the Priority Academic Program Development of Jiangsu Higher Education Institutions (PAPD), the 111 project (B16026), and China Scholarship Council, the Agriculture and Agri-Food Canada’s Project 1387 under the Canadian Agricultural Partnership, the DFG Research Unit FOR 1695 ‘Agricultural Landscapes under Global Climate Change – Processes and Feedbacks on a Regional Scale, the U.S. Department of Agriculture National Institute of Food and Agriculture (award no. 2015-68007-23133) and USDA/NIFA HATCH grant N. MCL02368, the National Key Research and Development Program of China (2016YFD0300105), The Broadacre Agriculture Initiative, a research partnership between University of Southern Queensland and the Queensland Department of Agriculture and Fisheries, the Academy of Finland through project AI-CropPro (315896), the JPI FACCE MACSUR2 project, funded by the Italian Ministry for Agricultural, Food, and Forestry Policies (D.M. 24064/7303/15 of 26/Nov/2015). The order in which the donors are listed is arbitrary.

