## Supplementary material for "How well do crop modeling groups predict wheat phenology, given calibration data from the target population?": Wallach_et_al_2020_SUPPLEMENTARY_MATERIAL

#### Running title: Crop model phenology prediction

Wallach[1], Daniel; Palosuo[2], Taru; Thorburn[3], Peter; Gourdain[4], Emmanuelle; Asseng[5], Senthold; Basso[6], Bruno; Buis[7], Samuel; Crout[8], Neil, Dibari[9], Camilla; Dumont[10], Benjamin; Ferrise[9], Roberto; Gaiser, Thomas[11]; Garcia[6], Cécile; Gayler[12], Sebastian; Ghahramani[13], Afshin; Hochman[3], Zvi; Hoek[14], Steven; Horan[3], Heidi; Hoogenboom[5,15], Gerrit; Huang[16], Mingxia; Jabloun[8], Mohamed; Jing[17], Qi; Justes[18], Eric; Kersebaum[19], Kurt Christian; Klosterhalfen[20], Anne; Launay[21], Marie; Luo[22], Qunying; Maestrini[6], Bernardo; Mielenz[23], Henrike; Moriondo[24], Marco; Nariman Zadeh[25], Hasti; Olesen[26], Jørgen Eivind; Poyda[27], Arne; Priesack[28], Eckart; Pullens[26], Johannes Wilhelmus Maria; Qian[17], Budong; Schütze[29], Niels; Shelia[5,15], Vakhtang; Souissi[30,31], Amir; Specka[19], Xenia; Srivastava[11], Amit Kumar; Stella[19], Tommaso; Streck[12], Thilo; Trombi[9], Giacomo; Wallor[19], Evelyn; Wang[16], Jing; Weber[12], Tobias, K.D.; Weihermüller[20], Lutz; de Wit[14], Allard; Wöhling[29,32], Thomas; Xiao[5,33], Liujun; Zhao[5], Chuang; Zhu[33], Yan, Seidel, Sabine J.[11]

[1]INRA, UMR AGIR, Castanet Tolosan, France. ORCID 0000-0003-3500-8179

[2]Natural Resources Institute Finland (Luke), Helsinki, Finland

[3]CSIRO Agriculture and Food, Brisbane, Queensland, Australia

[4]ARVALIS - Institut du végétal Paris, France

[5]Agricultural and Biological Engineering Department, University of Florida, Gainesville, Florida

[6]Department of Earth and Environmental Sciences, Michigan State University, East Lansing, Michigan

[7]INRA, UMR 1114 EMMAH, Avignon, France

[8]School of Biosciences, University of Nottingham, Loughborough, UK

[9]Department of Agriculture, Food, Environment and Forestry (DAGRI), University of Florence, Italy

[10]Department Terra & AgroBioChem, Gembloux Agro-Bio Tech, University of Liege, Gembloux, Belgium

[11]Institute of Crop Science and Resource Conservation, University of Bonn, Germany

[12]Institute of Soil Science and Land Evaluation, Biogeophysics, University of Hohenheim, Stuttgart, Germany

[13]Centre for Sustainable Agricultural Systems, Institute for Life Sciences and the Environment, University of Southern Queensland, Toowoomba, Queensland, Australia

[14] Environmental Sciences Group, Wageningen University & Research, Wageningen, The Netherlands
[15]Institute for Sustainable Food Systems, University of Florida, Gainesville, Florida
[16]College of Resources and Environmental Sciences, China Agricultural University, Beijing, China
[17]Ottawa Research and Development Centre, Agriculture and Agri-Food Canada, Ottawa, Canada
[18]CIRAD, UMR SYSTEM, Montpellier, France
[19]Leibniz Centre for Agricultural Landscape Research, Müncheberg, Germany
[20]Institute of Bio- and Geosciences - IBG-3, Agrosphere, Forschungszentrum Jülich GmbH, Jülich, Germany
[21]INRA, US 1116 AgroClim, Avignon, France
[22]Hillridge Technology Pty Ltd, Sydney, Australia
[23]Institute for Crop and Soil Science, Federal Research Centre for cultivated Plants, Julius Kühn-Institut (JKI), Braunschweig, Germany
[24]CNR-IBIMET, Firenze, Italy
[25]Aalto University School of Science, Espoo, Finland
[26]Department of Agroecology, Aarhus University, Tjele, Denmark
[27]Grass and Forage Science / Organic Agriculture, Institute of Crop Science and Plant Breeding, Kiel University, Kiel, Germany
[28]Institute of Biochemical Plant Pathology, Helmholtz Zentrum München-German Research Center for Environmental Health, Neuherberg, Germany
[29]Institute of Hydrology and Meteorology, Chair of Hydrology, Technische Universität Dresden, Dresden, Germany
[30]National Institute of Agronomic Research of Tunisia (INRAT), Agronomy Laboratory, University of Carthage, Tunis, Tunisia
[31]National Agronomy Institute of Tunisia (INAT), University of Carthage, Tunis, Tunisia
[32]Lincoln Agritech Ltd., Hamilton, New Zealand
[33]National Engineering and Technology Center for Information Agriculture, Jiangsu Key Laboratory for Information Agriculture, Jiangsu Collaborative Innovation Center for Modern Crop Production, Nanjing Agricultural University, Nanjing, Jiangsu, China

### SUPPLEMENTARY MATERIAL

**Table S1**

**Participating models, model version and references.**

| model | Version(s) | reference |
|---|---|---|
| AgroC | May2018 | Herbst M., Hellebrand H.J. , Bauer J., Huisman J.A., Šimůnek J., Weihermüller L., Graf A., Vanderborght J., Vereecken H. (2008). Multiyear heterotrophic soil respiration: Evaluation of a coupled CO2 transport and carbon turnover model. Ecological Modelling. 214: 271-283.<br><br>Klosterhalfen, A., Herbst M., Weihermüller L., Graf A., Schmidt M., Stadler A., Schneider K., Subke J.-A., Huisman J.A., Vereecken H. (2017). Multi-site calibration and validation of a net ecosystem carbon exchange model for croplands. Ecological Modelling. 363: 137-156. |
| APSIM | 7.8, 7.9, 7.10 | Keating B.A., P.S. Carberry, G.L. Hammer, M.E. Probert, M.J. Robertson, D. Holzworth, N.I. Huth, J.N.G. Hargreaves, H. Meinke, Z. Hochman, G. McLean, K. Verburg, V. Snow, J.P. Dimes, M. Silburn, E. Wang, S. Brown, K.L. Bristow, S. Asseng, S. Chapman, |

| | | |
|---|---|---|
| | | R.L. McCown, D.M. Freebairn and C.J.Smith. (2003). An overview of APSIM, a model designed for farming systems simulation. *European Journal of Agronomy 18: 267-288.*<br><br>Holzworth D.P., Huth N.I., DeVoil P.G. et al. (2014) APSIM - Evolution towards a new generation of agricultural systems simulation. Environmental Modelling & Software, 62, 327-350 |
| AquaCrop | 4.0 | Vanuytrecht E., Raes D., Steduto P., Hsiao T.C., Fereres E., Heng L.K., Garcia Vila M., Mejias Moreno, P. (2014). AquaCrop: FAO'S crop water productivity and yield response model. Environmental Modelling & Software, 62: 351-360 |
| CERES-Wheat | DSSATV4.7,V4.7., Expert-N 3.0 | Jones J.W., Hoogenboom G., Porter C.H., Boote K.J., Batchelor W.D., Hunt L.A., Wilkens P.W., Singh U., Gijsman A.J., Ritchie J.T. (2003). DSSAT Cropping System Model. European Journal of Agronomy 18, 235-265.<br><br>Hoogenboom G., Porter C. H., Shelia V., Boote K. J., Singh U., White J. W., Hunt L. A., Ogoshi R., Lizaso J. I., Koo J., Asseng S., Singels A., L.P. Moreno, Jones J. W. (2017). Decision Support System For Agrotechnology Transfer (DSSAT). Version 4.7. DSSAT |

| | | |
|---|---|---|
| | | Foundation, Gainesville, Florida, USA. |
| CROPSIM-Wheat | DSSAT V4.7 | Hoogenboom G., Porter C. H., Shelia V., Boote K. J., Singh U., White J. W., Hunt L. A., Ogoshi R., Lizaso J. I., Koo J., Asseng S., Singels A., L.P. Moreno, Jones J. W. (2017). Decision Support System For Agrotechnology Transfer (DSSAT). Version 4.7. DSSAT Foundation, Gainesville, Florida, USA. |
| Cropsyst | 3.04.08 | Stöckle C. O., Donatelli M., Nelson R. (2003). CropSyst, a cropping systems simulation model. European Journal of Agronomy, 18(3-4), 289-307. |
| DAISY | 5.59 | Hansen S., P. Abrahamsen C. T. Petersen, Styczen M.. (2012). Daisy: Model Use, Calibration, and Validation. Transactions of the ASABE, 55, 1317–1335. |
| Nwheat | DSSAT | Kassie B.T., Asseng S., Porter C.H. and Royce F.S. (2016). Performance of DSSAT-Nwheat across a wide range of current and future growing conditions. European Journal of Agronomy, 81, 27-36. |
| GECROS | Expert-N 3.0 | Yin X., van Laar H. H. (2005). Crop systems dynamics. An ecophysiological simulation |

| | | |
|---|---|---|
| | | model for genotype-by-environment interactions. Wageningen Academic Publishers, 155 pp., Wageningen, The Netherlands. |
| HERMES | 4.27 | Kersebaum K.C. (2007). Modelling nitrogen dynamics in soil-crop systems with HERMES. Nutrient Cycling in Agroecosystems, 77, 39-52.<br><br>Kersebaum K.C. (2011). Special features of the HERMES model and additional procedures for parameterization, calibration, validation, and applications In: L.R. Ahuja and L. Ma (ed.): Advances in Agricultural Systems Modeling Series 2. 65-94. ASA, CSSA, SSSA, Madison, USA. |
| LINTUL | LINTUL5 | Wolf J. (2012). User guide for LINTUL5: Simple generic model for simulation of crop growth under potential, water limited and nitrogen, phosphorus and potassium limited conditions. Wageningen UR. |
| MONICA | 2.02 | Nendel C., Berg M., Kersebaum K.C., Mirschel W., Specka X., Wegehenkel M., Wenkel K.O., Wieland R. (2011). The MONICA model: Testing predictability for crop growth, soil moisture and nitrogen dynamics. Ecological Modelling 222(9), 1614 - 1625. |

| | | |
|---|---|---|
| | | Specka X., Nendel C., Wieland R. (2015). Analysing the parameter sensitivity of the agro-ecosystem model MONICA for different crops. European Journal of Agronomy, 71, 73-87.<br><br>Specka X., Nendel C., Wieland R. (2019). Temporal Sensitivity Analysis of the MONICA Model: Application of Two Global Approaches to Analyze the Dynamics of Parameter Sensitivity. Agriculture 9(2), 37. |
| OpenCrop | | OpenCrop: An Open Source Crop Model – Model Description. Crout NMJ, Karanaratne, A & Jabloun, M (2018). School of Biosciences, University of Nottingham, UK |
| Salus | | Basso B, Ritchie JT, Grace PR, Sartori L (2006) Simulation of tillage systems impact on soil biophysical properties using the SALUS model. Italian Journal of Agronomy,1, 677-688.<br><br>Basso B. and J.T. Ritchie. 2015. Simulating Crop Growth and Biogeochemical Fluxes in Response to Land Management using the SALUS Model. In S. K. Hamilton, J. E. Doll, and G. P. Robertson, editors. The ecology of agricultural landscapes: long-term research on the path to sustainability. Oxford University Press, New York, NY USA |

| | | |
|---|---|---|
| SPASS | Expert-N 3.0 | Wang, E. (1997). Development of a Generic Process-Oriented Model for Simulation of Crop Growth. München, Herbert Utz Verlag Wissenschaft. 195 pp. |
| SSM-Wheat | | Soltani A., Maddah V., Sinclair T. (2013). SSM-Wheat: a simulation model for wheat development, growth and yield. International Journal of Plant Production, 7, 711-740. |
| STICS | 8_5_0 | Brisson N., Launay M., Mary B., Beaudoin N. (2009). Conceptual basis, formalisations and parametrization of the STICS crop model. Quae, 304pp<br><br>Coucheney E., Buis S., Launay M. Constantin J., Mary B., Garcia de Cortazar-Atauri I., Ripoche D., Beaudoin N., Ruget F., Andrianorisoa S., Le Bas C., Justes E., Léonard J. (2015). Accuracy, robustness and behavior of the STICS 8.2.2 soil-crop model for plant, water and nitrogen outputs: evaluation over a wide range of agro-environmental conditions in France. Environmental Modelling & Software, 64, 177-190 |
| XN-SUCROS | Expert-N 3.0 | van Laar, H.H. , J. Goudriaan, und H. van Keulen, 1992: Simulation of crop growth for potential and water-limited production situations (as applied to spring wheat).: Simulation Report CABO-TT no. 27. Wageningen: Centre for Agrobiological Research and Department |

| | | |
|---|---|---|
| | | of Theoretical Production Ecology, Wageningen Agricultural University;<br><br>Vanclooster, M., Viaene P., Diels J., Christiaens K., 1994: WAVE a mathematical model for simulating water and agrochemicals in the soil and vadose environment. Reference and user's manual (release 2.0). Leuven: Institute for Land and Water Management, Katholieke Universiteit Leuven. |
| PCWOFOST | 5.3.3 | Ceglar A. , van der Wijngaart R. , de Wit A., Lecerf R., Boogaard H., Seguini L. , van den Berg M., Toreti A., Zampieri M., Fumagalli D., Baruth B. (2019). Improving WOFOST model to simulate winter wheat phenology in Europe: Evaluation and effects on yield. Agricultural Systems. 168, 168-180. |
| WCCWOFOST | 7.1.7 | Boogaard, H.L., Van Diepen, C.A., Rötter, R.P., Cabrera, J.M.C.A., Van Laar, H.H., 1998. User's guide for the WOFOST 7.1 crop growth simulation model and WOFOST control center 1.5. Technical Document 52. Winand Staring Centre, Wageningen, the Netherlands, 144 pp. |
| Wheat-Grow | 3.1 | Zhu Y.; Liu L.; Liu, B. WheatGrow: A simulation model for predicting growth and |

| | | |
|---|---|---|
|  |  | productivity in wheat. In Proceedings of the Workshop on Modeling Wheat Response to High Temperature, Texcoco, Mexico, 19–21 June 2013.<br><br>Lv Z., Liu X., Tang L., Liu, L., Cao, W. and Zhu, Y., 2016. Estimation of ecotype-specific cultivar parameters in a wheat phenology model and uncertainty analysis. Agricultural and Forest Meteorology, 221: 219-229. |

.

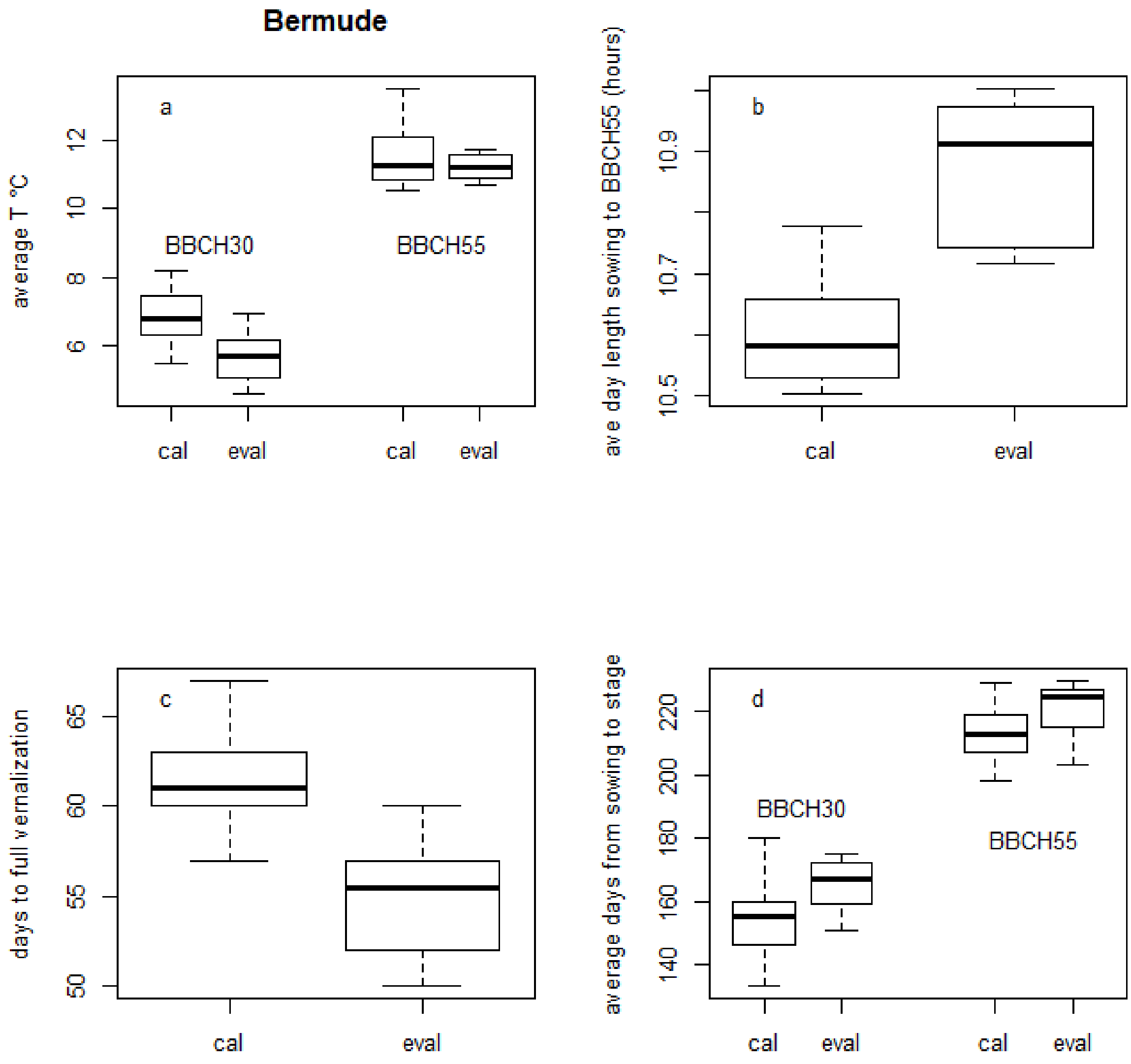


**Figure S1**

**Characteristics of calibration (“cal”) and evaluation (“eval”) data sets for variety Bermude. a) Boxplot of daily temperature averaged over the period from sowing to observed day of BBCH30 and from observed day of BBCH30 to observed day of BBCH55. b). Boxplot of day length (hours) averaged over the period from sowing to**

**observed day of BBCH55. c). Boxplot of days to full vernalization, calculated as explained in Materials and Methods d). Boxplot of observed days from sowing to BBCH30 and from sowing to BBCH55. The box encompasses 50% of the environments. The horizontal lone is the median. The ends of the whiskers show 1.5 times the interquartile range. Points outside that range are shown as circles.**

| | MAE eval | MSE eval | EF_eval | skillT eval | MAE cal |
|---|---|---|---|---|---|
| M17 | 3.47 | 19.78 | 0.88 | 0.88 | 3.77 |
| emedian | 3.53 | 21.09 | 0.86 | 0.88 | 3.68 |
| M15 | 3.72 | 22.72 | 0.86 | 0.86 | 3.09 |
| M2 | 3.94 | 27.13 | 0.83 | 0.85 | 3.80 |
| emean | 4.03 | 24.86 | 0.84 | 0.86 | 3.91 |
| M13 | 4.31 | 30.88 | 0.81 | 0.80 | 4.73 |
| M23 | 4.66 | 33.97 | 0.80 | 0.79 | 3.52 |
| M25 | 4.78 | 35.66 | 0.77 | 0.78 | 3.70 |
| M21 | 4.81 | 35.13 | 0.78 | 0.79 | 3.43 |
| M3 | 4.91 | 41.47 | 0.72 | 0.74 | 5.36 |
| M24 | 5.00 | 44.31 | 0.71 | 0.71 | 6.20 |
| M26 | 5.00 | 41.81 | 0.74 | 0.77 | 4.20 |
| M14 | 5.06 | 34.31 | 0.79 | 0.78 | 4.68 |
| M8 | 5.53 | 56.47 | 0.64 | 0.72 | 5.09 |
| M7 | 5.97 | 68.09 | 0.57 | 0.67 | 4.71 |
| M16 | 6.09 | 62.59 | 0.59 | 0.61 | 5.20 |
| M10 | 6.19 | 50.69 | 0.68 | 0.72 | 5.98 |
| M20 | 6.41 | 62.41 | 0.61 | 0.65 | 4.43 |
| M4 | 6.75 | 61.81 | 0.61 | 0.59 | 6.68 |
| M12 | 7.03 | 74.59 | 0.51 | 0.52 | 9.34 |
| M22 | 7.78 | 93.16 | 0.39 | 0.47 | 4.88 |
| M1 | 9.00 | 112.06 | 0.30 | 0.33 | 6.23 |
| M11 | 9.31 | 142.25 | 0.11 | 0.31 | 11.21 |
| M9 | 9.31 | 109.56 | 0.26 | 0.23 | 6.29 |

| | | | | | |
|---|---|---|---|---|---|
| M27 | 9.59 | 129.78 | 0.18 | 0.26 | 6.30 |
| M18 | 10.50 | 149.00 | -0.01 | -0.05 | 6.11 |
| M19 | 10.69 | 140.06 | 0.07 | 0.08 | 11.77 |
| onlyT | 11.03 | 172.47 | -0.09 | 0.00 | 8.07 |
| naive | 11.05 | 161.15 | 0.00 | 0.01 | 8.63 |
| M5 | 12.75 | 234.94 | -0.43 | -0.28 | 17.98 |
| M6 | 12.81 | 218.44 | -0.40 | -0.29 | 10.29 |

**Table S2.**

**Columns from left: MAE averaged over both stages and varieties, for the evaluation data (days), MSE averaged over both stages and varieties, for the evaluation data (days$^2$), EF averaged over both stages and varieties, for the evaluation data (unitless), skillT averaged over both stages and varieties, for the evaluation data unitless), and MAE averaged over both stages and varieties, for the calibration data (days). Models are in order of increasing MAE for the evaluation data.**
